# Approximately independent linkage disequilibrium blocks in human populations

**DOI:** 10.1101/020255

**Authors:** Tomaz Berisa, Joseph K. Pickrell

## Abstract

We present a method to identify approximately independent blocks of linkage disequilibrium (LD) in the human genome. These blocks enable automated analysis of multiple genome-wide association studies.

Availability (code) http://bitbucket.org/nygcresearch/ldetect

Availability (data): http://bitbucket.org/nygcresearch/ldetect-data

## 1 Introduction

The genome-wide association study (GWAS) is a commonly-used study design for the identification of genetic variants that influence complex traits. In this type of study, millions of genetic variants are genotyped on thousands to millions of individuals, and each variant is tested to see if an individual’s genotype is predictive of their phenotypes. Because of linkage disequilibrium (LD) in the genome [Pritchard and Przeworski, 2001], a single genetic variant with a causal effect on the phenotype leads to multiple statistical (but non-causal) associations at nearby variants. One initial analysis goal in a GWAS is to count the number of independent association signals in the genome while accounting for LD.

The most commonly-used approach to counting independent single nucleotide polymorphisms (SNPs) that influence a trait is to count “peaks” of association signals–this can be done manually when the number of peaks is small (e.g. Wellcome Trust Case Control Consortium [2007]), or in a semi-automated way when the number of peaks is larger (e.g. Jostins *et al.* [2012]). There are also fully automated methods that use LD patterns estimated from large reference panels of individuals [Yang *et al.*, 2012]. In some contexts (for example, when performing identical analysis on multiple GWAS with the goal of comparing phenotypes), it is useful to define approximately independent LD blocks *a priori* rather than letting them vary across analyses performed on different phenotypes [Loh *et al.*, 2015; Pickrell, 2014].

To define approximately-independent LD blocks, Loh *et al.* [2015] used non-overlapping segments of 1 megabase, and Pickrell [2014] used non-overlapping segments of 5,000 single nucleotide polymorphisms (SNPs). The breakpoints of these segments undoubtedly sometimes fall in regions of strong LD, thus potentially splitting a single association signal over two blocks (and leading to over-counting of the number of associated variants). A better approximation could be obtained by considering the empirical patterns of LD in a reference panel (e.g. Anderson and Novembre [2003]; Greenspan and Geiger [2004]; Mannila *et al*. [2003]). In the remainder of this paper, we present an efficient signal processing-based heuristic for choosing approximate segment boundaries.

## 2 Approach and Results

In order to estimate LD between pairs of SNPs, we use the *r*^2^ metric. If a genetic variant is in LD with another genetic variant that has a causal influence on disease, then *r*^2^ (times the strength of association at the causal SNP) is proportional to the association statistic at the non-causal SNP [Pritchard and Przeworski, 2001]. For our purposes, we define two sets of SNPs as “approximately independent” if the pairwise *r*^2^ between SNPs in different sets is close to zero.

Our approach is a heuristic for choosing segment boundaries, given a mean segment size (which is the required input). Let there be *n* genetic variants on a chromosome. The method can be broken down into the following basic steps (see the Supplementary Material for details):

1. Calculate the *n* × *n* covariance matrix *C* for all pairs of SNPs using the shrinkage estimator of *C* from Wen and Stephens [2010]
2. Convert the covariance matrix to *n* × *n* matrix of squared Pearson product-moment correlation coefficients *P*
3. Convert the matrix *P* = (*e*_*i,j*_) to a (2*n* − 1)-dimensional vector *V* = (*v*_*k*_) as follows:

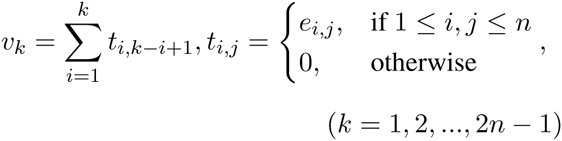 The effect of this step is representing each *antidiagonal* of *P* by the sum of its elements (Figs. 1a. and b.) This step has similarities to [Bulik-Sullivan *et al*., 2015], where the authors represent each *column* by the sum of its elements. The method presented in this paper uses the antidiagonal in order to differentiate between neighboring blocks of similar size.

**Figure 1:**
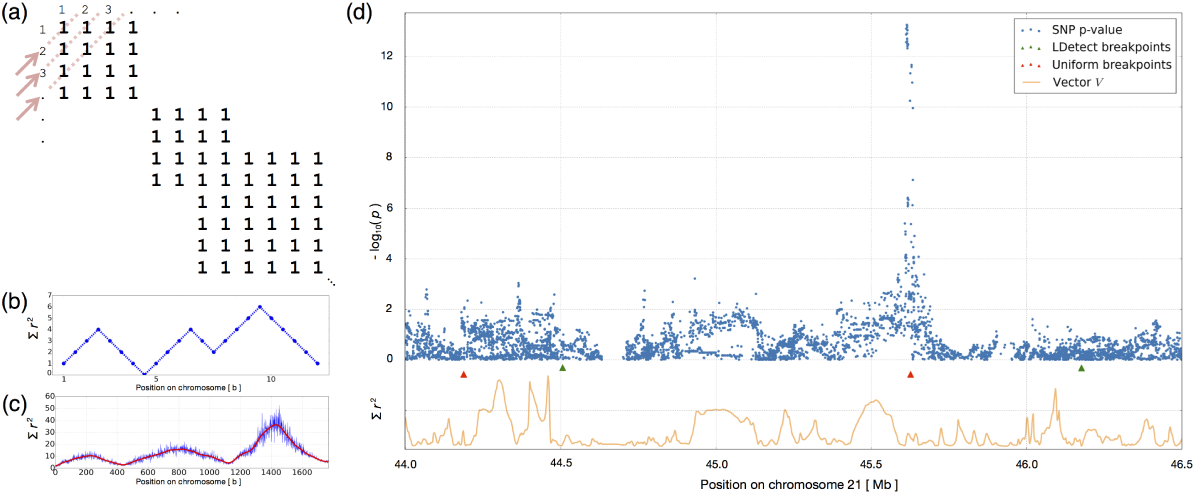
(a) and (b) Schematic of the conversion of matrix *P* to vector *V*. (c) Example data (blue) with Hann filter applied (red). (d) Example of Crohn’s disease GWAS hits with partially filtered vector *V* and comparison of breakpoints.
4. Apply low-pass filters of increasing widths to (i.e., *“smooth”*) *V* until the requested number of minima is achieved
5. Perform a local search in the proximity of each minimum from Step 4 in order to fine-tune the segment boundaries

In reality, matrix *P* turns out to be sparse, approx. banded, and approx. block-diagonal, with sporadically overlapping blocks [Slatkin, 2008; Wall and Pritchard, 2003; Wen and Stephens, 2010].

In order to provide intuition for Step 3, Fig. 1a. shows a simplified example of a correlation matrix *P*, where two SNPs *i* and *j* are either correlated (represented by 1 in element *e*_*ij*_ of the matrix) or uncorrelated (represented by zero, not shown). Representing each *antidiagonal* of *P* by the sum of its elements results in the vector shown in Fig. 1b. and identifying segments representing blocks of LD reduces to identifying local (or more stringently, global) minima in this vector. In reality, the elements *e*_*ij*_ of *P* are continuous values from the interval [0, 1] and result in an extremely noisy vector *V* (example in blue in Fig. 1c.) Therefore, in order to identify large-scale trends of LD and reduce high frequency components in the signal, we apply a signal processing technique dubbed low-pass filtering (utilizing a Hann window [Blackman and Tukey, 1958]) in Step 4. The result of applying a low-pass filter (with *width* = 100) is shown in red in Fig. 1c.

Applying wider and wider filters to vector *V* in Step 4 allows us to focus on the large scale structure of LD blocks, but also causes the approach to miss small scale variation around identified minima. In order to counteract this effect, Step 5 conducts a local search in the proximity of each local minimum identified in Step 4 to find the closest SNP *l* with 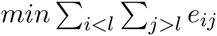.

We applied this method to sequencing data from European, African, and East Asian populations in the 1000 Genomes Phase 1 dataset. We set a mean block size of 10,000 SNPs, and used the algorithm to define the block boundaries. As expected, these boundaries fall in regions with considerably higher recombination rates than the genome-wide average (Supplementary Figure 4). In Fig. 1d., we show an example from genome-wide association study (GWAS) results for Crohn’s disease [Jostins *et al.*, 2012] where using uniformly-distributed breakpoints would result in double-counting of an association signal, while the LD-aware breakpoints avoid stretches of SNPs in LD.

To test whether this approach is useful more generally, we ran fgwas [Pickrell, 2014] on GWAS of Crohn’s disease [Jostins *et al.*, 2012] and height [Wood *et al.*, 2014], using both uniformly-distributed breakpoints and LD-aware breakpoints. Using the LD-aware breakpoints successfully eliminated double-counting of SNPs in moderate-to-high LD and on opposite sides of uniform breakpoints (Supplementary Materials, Section 6).

## References

Anderson, E. C. and Novembre, J. (2003). Finding haplotype block boundaries by using the minimum-description-length principle. Am J Hum Genet, 73(2), 336–54.

Blackman, R. B. and Tukey, J. W. (1958). The measurement of power spectra from the point of view of communications engineering - part i. Bell System Technical Journal, 37(1), 185–282.

Bulik-Sullivan, B. K., Loh, P.-R., Finucane, H. K., Ripke, S., Yang, J., Patterson, N., Daly, M. J., Price, A. L., Neale, B. M., of the Psychiatric Genomics Consortium, S. W. G., et al. (2015). Ld score regression distinguishes confounding from polygenicity in genome-wide association studies. Nature genetics, 47(3), 291–295.

Greenspan, G. and Geiger, D. (2004). Model-based inference of haplotype block variation. J Comput Biol, 11(2-3), 493–504.

Jostins, L., Ripke, S., Weersma, R. K., Duerr, R. H., McGovern, D. P., Hui, K. Y., Lee, J. C., Schumm, L. P., Sharma, Y., Anderson, C. A., et al. (2012). Host-microbe interactions have shaped the genetic architecture of inflammatory bowel disease. Nature, 491(7422), 119–124.

Loh, P.-R., Bhatia, G., Gusev, A., Finucane, H. K., Bulik-Sullivan, B. K., Pollack, S. J., de Candia, T. R., Lee, S. H., Wray, N. R., Kendler, K. S., et al. (2015). Contrasting regional architectures of schizophrenia and other complex diseases using fast variance components analysis. bioRxiv, page 016527.

Mannila, H., Koivisto, M., Perola, M., Varilo, T., Hennah, W., Ekelund, J., Lukk, M., Peltonen, L., and Ukkonen, E. (2003). Minimum description length block finder, a method to identify haplotype blocks and to compare the strength of block boundaries. Am J Hum Genet, 73(1), 86–94.

Pickrell, J. K. (2014). Joint analysis of functional genomic data and genome-wide association studies of 18 human traits. The American Journal of Human Genetics, 94(4), 559–573.

Pritchard, J. K. and Przeworski, M. (2001). Linkage disequilibrium in humans: models and data. The American Journal of Human Genetics, 69(1), 1–14.

Slatkin, M. (2008). Linkage disequilibrium?understanding the evolutionary past and mapping the medical future. Nature Reviews Genetics, 9(6), 477–485.

Wall, J. D. and Pritchard, J. K. (2003). Haplotype blocks and linkage disequilibrium in the human genome. Nature Reviews Genetics, 4(8), 587–597.

Wellcome Trust Case Control Consortium (2007). Genome-wide association study of 14,000 cases of seven common diseases and 3,000 shared controls. Nature, 447(7145), 661–78.

Wen, X. and Stephens, M. (2010). Using linear predictors to impute allele frequencies from summary or pooled genotype data. The annals of applied statistics, 4(3), 1158.

Wood, A. R., Esko, T., Yang, J., Vedantam, S., Pers, T. H., Gustafsson, S., Chu, A. Y., Estrada, K., Luan, J., Kutalik, Z., et al. (2014). Defining the role of common variation in the genomic and biological architecture of adult human height. Nature genetics, 46(11), 1173–1186.

Yang, J., Ferreira, T., Morris, A. P., Medland, S. E., Madden, P. A., Heath, A. C., Martin, N. G., Montgomery, G. W., Weedon, M. N., Loos, R. J., et al. (2012). Conditional and joint multiple-snp analysis of gwas summary statistics identifies additional variants influencing complex traits. Nature genetics, 44(4), 369–375.

